# GABA and astrocytic cholesterol determine the lipid environment of GABA_A_R in cultured cortical neurons

**DOI:** 10.1101/2024.04.26.591395

**Authors:** Zixuan Yuan, Mahmud Arif Pavel, Scott B. Hansen

## Abstract

The γ-aminobutyric acid (GABA) type A receptor (GABA_A_R), a GABA activated pentameric chloride channel, mediates fast inhibitory neurotransmission in the brain. The lipid environment is critical for GABA_A_R function. How lipids regulate the channel in the cell membrane is not fully understood. Here we employed super resolution imaging of lipids to demonstrate that the agonist GABA induces a rapid and reversible membrane translocation of GABA_A_R to phosphatidylinositol 4,5-bisphosphate (PIP_2_) clusters in mouse primary cortical neurons. This translocation relies on nanoscopic separation of PIP_2_ clusters and lipid rafts (cholesterol-dependent ganglioside clusters). In a resting state, the GABA_A_R associates with lipid rafts and this colocalization is enhanced by uptake of astrocytic secretions. These astrocytic secretions enhance endocytosis and delay desensitization. Our findings suggest intercellular signaling from astrocytes regulates GABA_A_R location based on lipid uptake in neurons. The findings have implications for treating mood disorders associated with altered neural excitability.

## Introduction

The γ-aminobutyric acid type A receptor (GABA_A_Rs) is an anesthetic channel essential for regulating neuronal excitability. This pentameric chloride channel, activated by the neurotransmitter γ-aminobutyric acid (GABA, Fig. S1a), initiates a sequence of events that leads to channel opening and influx of chloride ion (Fig. S1b). This influx results in the hyperpolarization of neurons, thereby reducing their firing rate.

Lipids, in particular cholesterol, plays an important regulatory role in GABA_A_R functionality, affecting its responsiveness to various modulators^1,2^. Notably, cholesterol enrichment diminishes the effectiveness of bicuculline, a competitive antagonist, while both enrichment and depletion alter responsiveness to GABA activation^3^. These cholesterol-induced changes, in conjunction with effects on other channels, significantly impact neuronal function and excitability^4^. The effects of cholesterol on GABA_A_R are thought to occur through both direct binding and indirect effects through the properties of the membrane^5,6^. Understanding cholesterol’s regulation of GABA_A_R is necessary for understanding cholesterol’s regulation of neuronal excitability.

Astrocytes, the brain’s primary cholesterol producer, secrete cholesterol and other minor lipids which are transported to neurons in lipid particles that contain apolipoprotein E (ApoE)^7,8^. The astrocyte-derived cholesterol, and associated lipids, are vital for functional integrity and nanoscopic spatial patterning of proteins within lipid compartments^8,9^ (Fig. S1c). Specifically, for GABA_A_R, cholesterol increases the channel’s association with lipid rafts in neurons^10^.

The localization of GABA_A_R within lipid rafts and within the synapse is dictated by the presence of the γ2 subunit ^11–14^. Within the synapse, GABA_A_R is further localized to gephyrin containing synaptic sub domains (SSDs) ^15–19^. The lipid composition of the SSDs is not well defined, but both GABA_A_R and gephyrin are palmitoylated (a saturated 16 carbon lipid covalently attached to the protein) and palmitoylation regulates raft affinity of most proteins^20^ and specifically enhances GABA_A_R’s presence within lipid rafts and with monosialotetrahexosylgangliosides (GM1)^20–22^, a common marker of lipid rafts^20,23,24^ (see Fig. S1).

GM1 lipids, identifiable by cholera toxin B (CTxB) labeling, are separate from lipid clusters containing phosphatidylinositol 4,5-bisphosphate (PIP_2_)^25^. PIP_2_ is an abundant anionic signaling lipid typically found on the inner leaflet of the plasma membrane. In untreated cultured cells, GM1 and PIP_2_ clusters are 50-200 nm apart^26,27^. This spatial separation allows for dynamic changes in a protein’s association with GM1 and PIP_2_ clusters and subsequent exposure to signaling lipids in the distinct compartments^9,28–31^. In contrast to the prototypical activation mechanism where the lipid ligand increases in concentration to activate the protein, here the protein moves to a specific lipid compartment which effectively exposes the protein to a high concentration of the lipid. Anesthetics facilitate this shift by competing with palmitate in lipid rafts (GM1 containing clusters)^32,33^ (Fig. S1d-e). The site thus referred to as an anesthetic/palmitate (AP) site^34^.

At its *α*1 subunit interface, GABA_A_R binds PIP_2_^35^. Unlike direct activation of potassium channels^36,37^, PIP_2_ binding to GABA_A_R is thought to affect receptor trafficking^35^. Here we demonstrate that at the nanoscopic level, the endogenous agonist GABA induces GABA_A_R to dissociate from GM1 clusters and associate with PIP_2_ cluster, a process regulated by astrocyte cholesterol. The astrocyte cholesterol opposes GABA-induced dissociation of GABA_A_R from lipid rafts and the movement coincides with a delay in receptor activation and desensitization. We propose a positional activation model wherein the lipids in neurons regulate the location of GABA_A_R, and astrocytes supply the lipids that regulate the channel. Hence astrocytes play a crucial role in neuronal excitability through brain derived lipids.

## Results

### Spatial Patterning of GABA_A_R and GM1 lipids in primary neurons through two-color dSTORM

To determine the spatial relationship of GABA_A_R with GM1 lipid clusters, we employed two-color direct stochastic optical reconstruction microscopy (dSTORM) to image fixed primary cortical neurons in a resting state from day 17 mouse embryos (Fig. 1a). These neurons naturally express GABA_A_R, GM1, and PIP_2_, allowing for the visualization of GM1 and PIP_2_ clusters, which are typically beyond the diffraction limit of conventional optical microscopy^38^. The fixation process, employing both paraformaldehyde and glutaraldehyde, was performed both before and after antibody labeling. This approach ensured that protein movement between lipid domains and potential changes in the structure of lipid domains themselves were minimized during imaging. The specificity of the antibodies used was confirmed by previous validation studies^39–41^. See also Fig.S2a-f and supplemental discussion.

**Figure 1.**
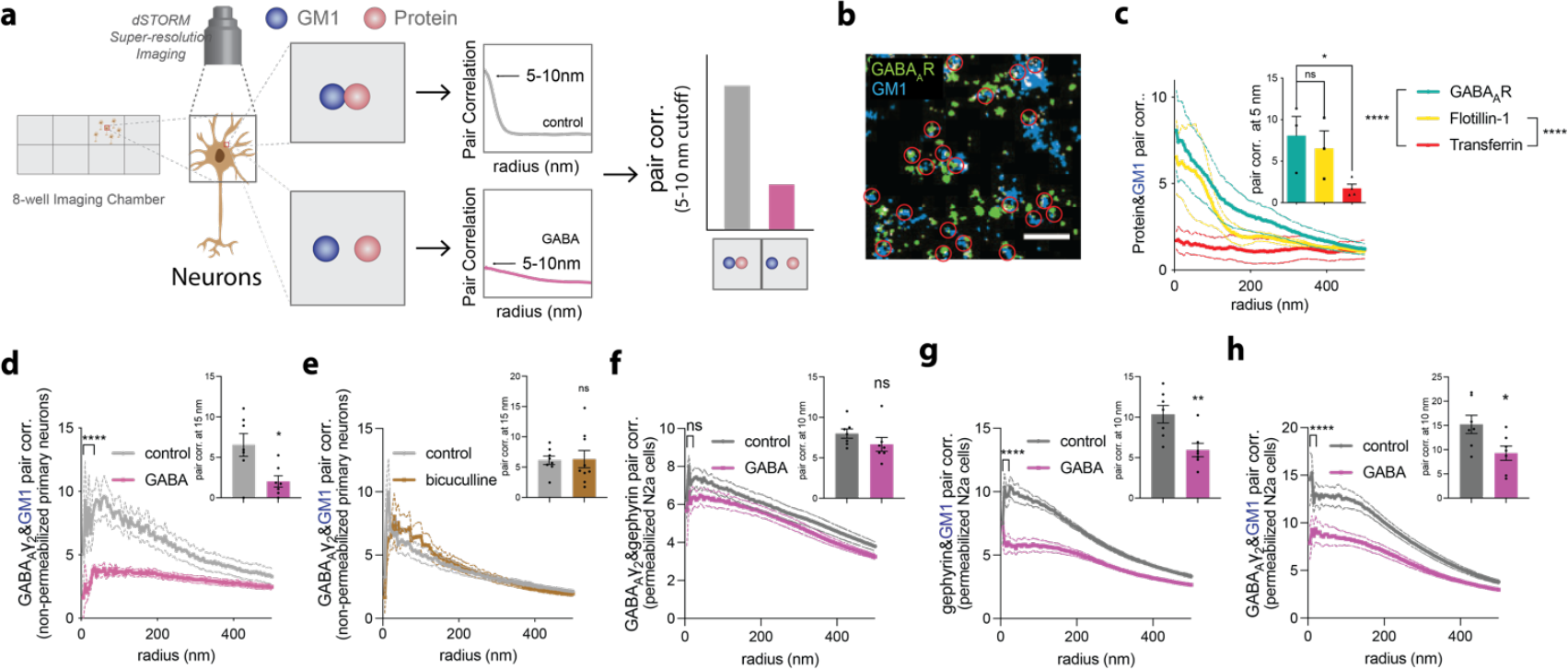
GABA agonism causes GABA_A_R to disassociate from GM1 clusters. **(a)** A schematic representation illustrates the experimental setup for two-color super-resolution imaging, depicting the association of gamma amino butyric acid (GABA) type A receptor (GABA_A_R) containing the gamma 2 subunit (GABA_A_γ2, red) with GM1 lipids (blue) and the methodology for analyzing spatial distribution through cross-pair correlation curves. The diagram indicates that a high cross-pair correlation (pair corr.) suggests association between GABA_A_R and GM1 lipids, which diminishes following treatment with GABA. **(b)** Representative dSTORM image of GABA_A_R and GM1 labeling in primary cortical neurons. Red circles indicate regions of GABA_A_R/GM1 colocalization. **(c)** Pair corr. of GABA_A_γ2, flotillin, transferrin and GM1 in primary neurons determined by two-color dSTORM imaging (n=3-4). Statistical comparison of pair corr. curves using a nested t-test at the radii 5-25 nm—results are shown next to the figure legend. A bar graph at the shortest distance, 5 nm, shows quantification of the change in pair corr. at a single point. **(d)** Pair corr. analysis of GABA_A_γ2 and GM1 dSTORM imaging in primary neurons treated with 1mM GABA (endogenous agonist) for 10 mins (n=7). The radius used for the nested t-test is 10-35 nm. Bar graph shows quantification of a single point at 15 nm. **(e)** Pair corr. of GABA_A_γ2 and GM1 in primary neurons treated with 10 μM bicuculline (GABA_A_R antagonist) for 10 mins (n=9-10). Bar graph shows quantification of the curve at 15 nm. **(f)** Pair corr. analysis of GABA_A_γ2 and gephyrin in N2a cells treated with 1 mM GABA for 10min (n=7). The radius used for the nested t-test is 5-25 nm. The bar graph shows quantification at 10 nm radius. **(g)** Pair corr. analysis of gephyrin and GM1 in N2a cells treated with 1 mM GABA for 10min (n=7). The two curves were compared with a nested t-test using radii of 5-25 nm. The bar graph shows quantification at 10 nm radius. **(h)** Pair corr. analysis of GABA_A_γ2 and GM1 in N2a cells treated with and without 1 mM GABA for 10min (n=7). A nested t-test using radii from 5-25 nm was used to compare the two curves. The bar graph shows quantification at 10 nm radius. All data are expressed as mean ± s.e.m., ns, not significant, *p<0.05, **p<0.01, ****p<0.0001, unless otherwise noted. Each n is a biological replicate.

A representative dSTORM image (Fig. 1b) shows the distribution and co-localization of GABA_A_R (green) and GM1 lipid clusters (blue) in cultured primary neurons. The image is a statistical composite derived from 1000 to 3000 individual images, totaling approximately 150,000 particles for each probe, captured at 60x magnification with a 100 ms exposure per frame. The GM1 clusters we describe here are the same as the lipid rafts obtained from detergent resistant membranes, however when referring to our measurements, we use the term “GM1 clusters” or “GM1 domains”, since that is what we label and measure.

Cross-pair correlation (pair corr.) analysis of single molecule localizations (SML) produced a curve indicating the proximity of GABA_A_R to GM1 domains. The curve’s absolute reading can vary based on different cell types or labeling methods; therefore, treatments were performed in parallel and compared directly to their own control. From our observations in mouse primary cortical neurons and neuroblastoma 2a (N2a) cells, the curve’s unitless value of >5 near the y axis suggest a significant association between GABA_A_R and GM1 (Fig. 1c). Values of 2-3 are considered weak or no correlation and value less than 1 are anti-correlated (see methods). The cross-pair correlation can also be seen in Fig. 1c inset showing a single point at the shortest distance (5 nm) along the curve.

To confirm GABA_A_R association with lipid rafts, we compared GABA_A_R/GM1 cross-pair correlation with flotillin/GM1 cross-pair correlation, the latter is a known protein marker for lipid rafts ^42^ (Fig.1c, S2g). The comparison showed that GABA_A_R/GM1 association was like that of flotillin/GM1, while transferrin, a non-raft protein^43,44^, showed negligible association with GM1 clusters (Fig.1c, S2h). The difference in cross-pair correlation at a 5 nm radius among GABA_A_R/GM1, flotillin/GM1, and transferrin/GM1 is highlighted in the inset of Fig. 1c. Again, high cross-pair correlation values at this short distance indicate a strong association with GM1 domains, with GABA_A_R and flotillin showing similar patterns, contrasting significantly with transferrin. These findings collectively suggest that GABA_A_R is localized to GM1 containing lipid domains within the neuronal cell membrane.

### Effects of GABA activation on GABA_A_R and lipid localization

Next, we explored how GABA_A_R localization changes in response to GABA, the receptor’s endogenous agonist. We hypothesized that GABA activation could lead to a reorganization of GABA_A_R’s spatial patterning, correlating a physiological condition with the patterning. To this end, primary cortical neurons were treated with 1 mM GABA for 10 minutes, fixed, stained (see Fig. S2a-c for representative images), analyzed via two-color dSTORM, and changes in localization compared to untreated controls. The 1 mM concentration of GABA was decided based on the concentration present in human brain and a maximum activation effect reported in the literature^45–48^. And 10 min was used to assure that the receptor had reached its final state.

GABA treatment resulted in a significant reduction in the association of GABA_A_R with GM1 clusters, decreasing by more than 56% ± 13% (Fig. 1d, p<0.001). The pattern was corroborated in cultured neuroblastoma 2a (N2a) cells, which also stably express GABA_A_Rγ2^49^, displaying similar decreases in GABA_A_R/GM1 association (Fig. 2a). Conversely, treatment with bicuculline, a GABA_A_R competitive antagonist that targets the same site as GABA^50,51^, did not affect the receptor’s localization (Fig. 1e). Importantly, GABA did not disrupt lipid rafts, as evidenced by unchanged activity of phospholipase D (PLD), a raft dependent protein, and cholesterol levels following treatment (Fig. S3a-c and Fig. S3d-f, respectively).

**Figure 2.**
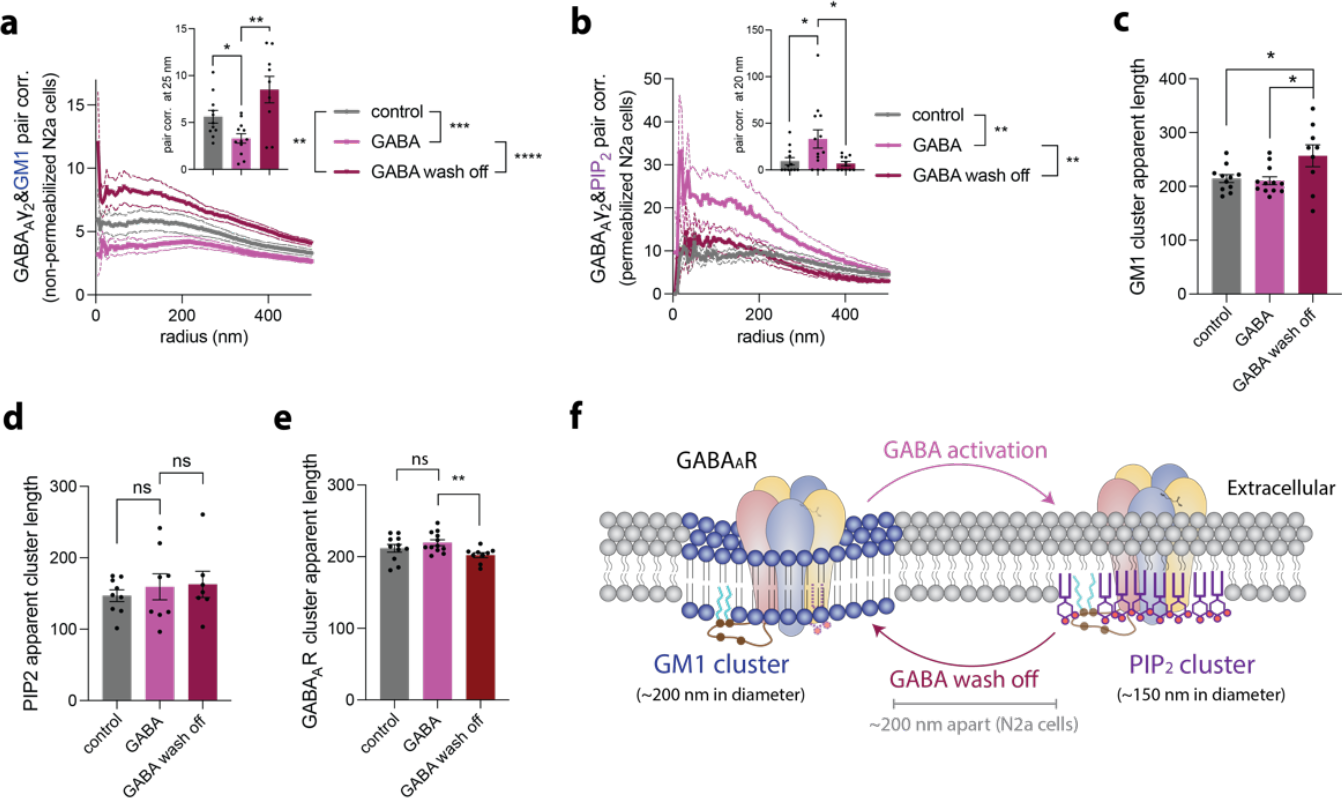
GABA treatment induces a reversible association of GABA_A_R with PIP_2_ clusters. **(a)** Cross-pair correlation (pair corr.) analysis of GABA_A_γ2 and GM1 in N2a cells treated with 1 mM GABA (endogenous agonist) for 10min and cells that are washed off for 15min after 10min 1 mM GABA treatment (n=9-12). The curves were compared using a nested t-test using radii from 5-30 nm. The bar graph shows quantification at a 25 nm radius. **(b)** Pair corr. analysis of GABA_A_γ2 and PIP_2_ in N2a cells treated with 1 mM GABA for 10 min. For GABA wash off, the 1 mM GABA was removed after 10 min and allowed to recover for 15 min (n=12-13). The radius used for nested t-test is 10-35 nm. The bar graph show quantification at 20 nm radius. **(c)** Apparent length of GM1 clusters in non-permeabilized N2a cells (n=9-12). **(d)** Apparent length of PIP_2_ clusters in permeabilized N2a cells (n=7-9). **(e)** Apparent length of GABA_A_R clusters in non-permeabilized N2a cells(n=9-12). **(f)** A cartoon diagram of GABA_A_R translocation from GM1 to PIP_2_ clusters upon GABA activation. The process is revered upon GABA removal. All data are expressed as mean ± s.e.m. using a t-test; ns, not significant, *p<0.05, **p<0.01, ***p<0.001, ****p<0.0001. Each n is a biological replicate.

To further confirm a translocation associated with GABA_A_R activation, we investigated the spatial relationship between GABA_A_R and gephyrin, a peripheral membrane protein that forms a complex with GABA_A_R, using three-color dSTORM (Fig. 1f-h). We reasoned gephyrin may also change its association with lipid rafts if the complex remained intact after GABA treatment. To this end we co-labeled GABA_A_R, gephyrin, and GM1 clusters on N2a cells. The cells were permeabilized for intracellular accessibility of antibody.

We found that GABA treatment led to a minor, non-statistically significant decrease in the cross-pair correlation between GABA_A_R and gephyrin (Fig. 1f). Moreover, the treatment caused a significant ~43% reduction in gephyrin cross-pair correlation with GM1 lipids (Fig. 1g), coupled with a corresponding ~39% decrease in the cross-pair correlation between GABA_A_R and GM1 clusters within the same cells (Fig. 1h). These data suggest that after GABA activation of GABA_A_R, at least a portion of the complex appears to remain intact and changes its position relative to GM1 lipid rafts.

Notably, when co-treating the cells with GABA combined with the antagonist bicuculline, a high concentration of bicuculline (100 μM) reversed the effect of 1mM GABA shifting the receptor out of GM1, whereas a low concentration of bicuculline (10 μM) did not cause this reversal (Fig. S3l). This result suggests a direct effect through GABA_A_R but does not by itself exclude GABA signaling through the type B metabotropic GABA receptor (GABA_B_R).

In addition to activating GABA_A_R, GABA binds to and activates GABA_B_R, and this activation enhances GABA_A_R current^52^. To investigate a potential contribution of GABA_B_R to the observed change in GABA_A_R localization, we treated N2a cells with the GABA_B_R antagonist saclofen, and determined the cross-pair correlation of natively expressed GABA_A_R and GM1 after 1 mM GABA treatment, identical to our bicuculline experiment. The expression of GABA_B_R has been shown in other neuroblastoma cell lines^53^.

After 10 min exposure to 100 μM saclofen, we saw no significant change in the cross-pair correlation of GABA_A_R with GM1 lipids (Fig. S3m). And unlike bicuculline, high concentration of saclofen failed to block GABA (1 mM) induced translocation of GABA_A_R. Overall, these data suggest that the effect of GABA on GABA_A_R translocational activation out of GM1 domains appears primarily direct through an action on GABA_A_R and not indirect through the activation of the GABA_B_R pathway.

### Reversible colocalization of GABA_A_R with PIP_2_ post-GABA treatment

Next, we investigated the fate of GABA_A_R after dissociation with raft lipids. As mentioned GABA_A_R was recently shown to bind to PIP_2_^35^. To assess GABA_A_R’s association with PIP_2_, a lipid primarily located on the plasma membrane’s inner leaflet, we treated N2a cells for 10 min with 1 mM GABA, permeabilized the cell membrane, and immuno-stained for PIP_2_ and GABA_A_R (Fig. S2d-f). Prior to GABA_A_R treatment, GABA_A_R/PIP_2_ cross-pair correlation analysis showed low cross-pair correlation values. However, post-GABA treatment, this association increased dramatically (256% ± 116%, Fig. 2b), suggesting that GABA_A_R dissociates from GM1 and associates with PIP_2_ in response to GABA. GABA treatment had minimal effect on GABA_A_R and PIP2 cluster number and size (Fig. S3h-k).

Many native biological processes are reversible, including GABA_A_R activation. Establishing reversibility is helpful for establishing a physiologically relevant process. To ascertain the reversibility of this GABA-induced relocation to PIP_2_ clusters, cells initially treated with GABA (1 mM for 10 min) were washed and incubated in fresh buffer (15 min) before fixation. This procedure revealed that GABA_A_R’s association with PIP_2_ domains is indeed reversible; washing out GABA resulted in a significant decrease in PIP_2_ association and a concomitant increase in GM1 association (Fig. 2a-b). These findings confirm the dynamic and reversible nature of GABA_A_R localization in response to GABA agonism.

### Molecular model of GABA_A_R localization dynamics

Figure 2f proposes a molecular model illustrating GABA_A_R’s dynamic regulation by lipid localization. Initially, receptors are associated with GM1 clusters. Upon activation by GABA, they transition to PIP_2_ clusters, and this relocation is reversed upon the removal of GABA. Measurements in N2a cells show GM1 cluster diameter average 215 nm, PIP_2_ clusters 147 nm, with an average separation of 207 nm (Fig. 2c-d, S1f).

### Latency of GABA induced GABA_A_R dissociation from GM1 clusters in live cells

Understanding the timing and latency of neuronal processes is crucial for delineating their potential physiological roles in the brain. To investigate the temporal dynamics of GABA-induced dissociation of GABA_A_R from GM1 clusters, we employed live-cell two-color dSTORM of the receptor with lipid. We optimized the labeling protocol using a 30-minute staining duration, followed by three brief washes immediately before imaging without fixation, ensuring effective labeling of both lipids and proteins (Fig. 3a).

**Figure 3.**
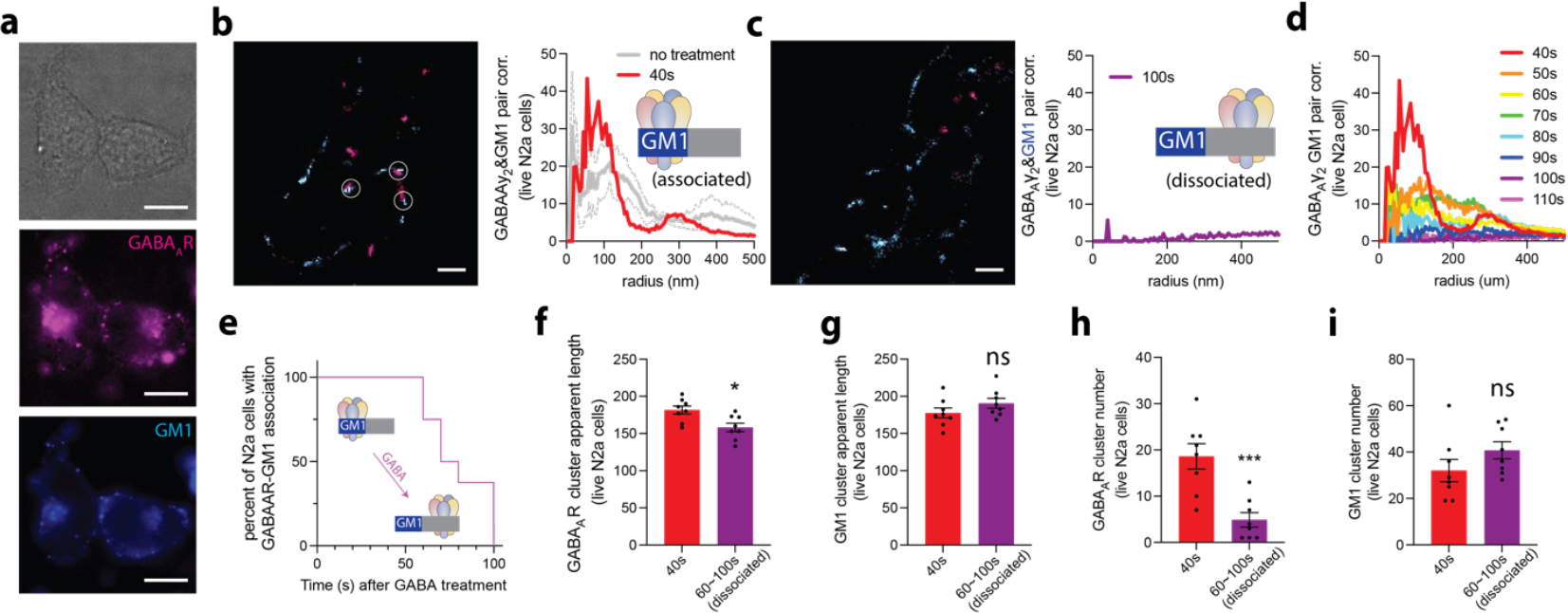
Live-cell imaging of GABA_A_R moving between lipid membrane compartments. **(a)** Representative images showing fluorescent labeling of lipids and GABA_A_R in live cells. **(b)** (left) A representative image of a two-color dSTORM image in a live N2a cell. The association of GABA_A_R and GM1 is pointed out by white circles. (right) Cross-pair correlation (pair corr.) analysis of no treatment and the shortest time point (40 s) after 1 mM GABA treatment. **(c)** (left) A representative dSTORM image of the same cell in (b) after 100s—there are no overlapping regions to label. (right) Pair corr. analysis of the data at 100 s (purple line). The cartoon depicts GABA_A_R dissociated from GM1 lipids (blue shading) and associated with disordered lipids (grey shading). **(d)** A time course of the GABA_A_R/GM1 Pair corr. from the cell in panel (b) after treatment with 1mM GABA. **(e)** Survival curve showing the percent of cells with pair corr. after 1 mM GABA treatment. GABA_A_R fully dissociated from GM1 lipids in all cells by 100s. **(f)** Apparent length of GABA_A_R clusters in live N2a cells (n=8). **(g)** Apparent length of GM1 clusters in live N2a cells (n=8). **(h)** Number of GABA_A_R clusters measured in live N2a cells (n=8). **(i)** Number of GM1 clusters in live N2a cells (n=8). All data are expressed as mean ± s.e.m., unpaired t-test, ns, not significant, *p<0.05, ***p<0.001. Each n value is a biological replicate.

The exposure time was set to 5 ms. Data collection for GABA_A_R utilized 800 images over 4 seconds (~12,500 particles), while GM1 clusters were captured in 200 images over 1 second (~30,000 particles). This setup allowed for the calculation of two-color cross-pair correlation analysis every 5 seconds, with data analysis binned at 10-second intervals. The cells were first located and focused under the microscope in 50μL imaging buffer. Subsequently, another 50μL imaging buffer containing 2mM GABA was added to achieve the final 1mM concentration, followed by drug equilibration. Due to the time needed for the safety latch to reset (~40 seconds), we used 1 mM GABA, which is a high physiological concentration and results in a slow desensitization rate of the receptor^54–57^.

Comparative analysis of live two-color SML showed a marked decrease in the cross-pair correlation between GABA_A_R and GM1 after 1 mM GABA treatment. In Figure 3b-d we present one cell as an example. Initially, after adding GABA to the imaging chamber, GABA_A_R clusters were observed near GM1 clusters (as indicated by white circles in Fig. 3b, left panel, and cross-pair correlation curve in Fig. 3b, right panel). Following GABA treatment, this association diminished (Fig. 3c). Figure 3d shows the whole dissociation process in this example cell. Averaging multiple cells, we found GABA_A_R requires an average of almost 80 seconds to fully dissociate from GM1 clusters in live cells (n=8) (Fig. 3e). In contrast, without any treatment, the cross-pair correlation between GABA_A_R and GM1 clusters in live N2a cells remained unchanged at the initial time point (Fig. 3b). All cells were fully dissociated by 100 s (~ 2 min.) confirming our 10 min incubation with GABA in fixed cells had reached a steady state.

Cluster analysis in live cells revealed no significant change in GM1 lipid cluster diameter or number following GABA treatment, but a decrease in GABA_A_R cluster diameter (182 nm to 158 nm) (Fig. 3f-i), consistent with a move of the receptor towards smaller PIP_2_ clusters (Fig. 2c-d). This shift indicates a direct correlation between receptor localization and functional state changes, underscoring the intricate balance between lipid environments and receptor dynamics.

### Astrocytic regulation of GABA_A_R localization in cultured neurons

#### GABA_A_R and GM1 clusters

Cholesterol’s role in regulating GM1 clustering highlights its significance in cellular signaling and spatial patterning. In the brain, cholesterol is predominantly synthesized by astrocytes and then transported to neurons with apolipoprotein E (ApoE) containing particles^8,9^. To investigate the impact of inter-cellular cholesterol derived from astrocytes on the spatial patterning of GABA_A_R, we utilized astrocyte-conditioned media (ACM) to treat primary cultured neurons and neuroblastoma 2a (N2a) cells. ACM was prepared by culturing primary cortical astrocytes in 10% FBS DMEM and collecting the media after 3 days, which contains cholesterol among other components^9^ (Fig. 4a).

**Figure 4.**
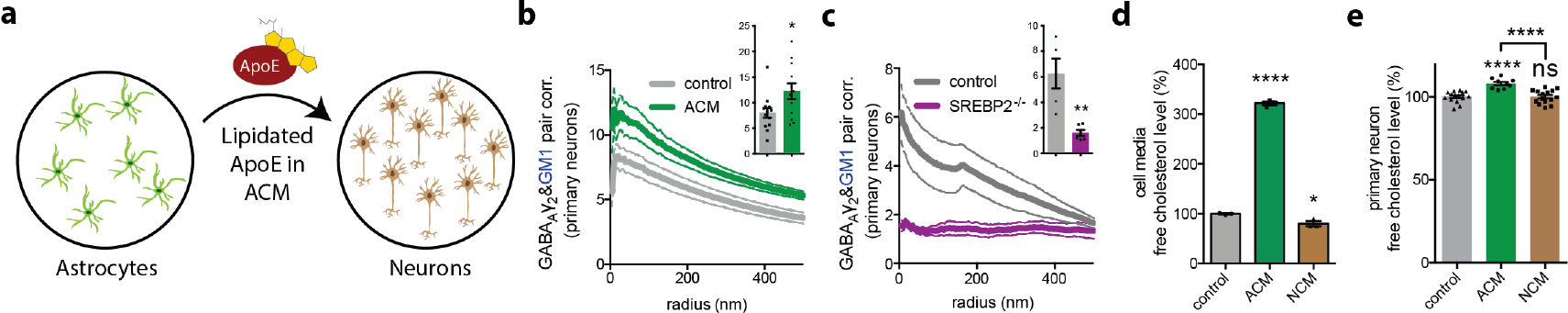
Astrocyte conditioned media drives GABA_A_R clustering with GM1 in neurons. **(a)** Cartoon showing the experimental setup. The astrocyte conditioned media (ACM) containing lipid secretions are collected from astrocyte culture and transferred to neuronal culture where it is taken up by the neuron in the presence of apolipoprotein E (ApoE). **(b)** GABAAγ2/GM1 pair corr. analysis in primary neurons treated with and without ACM (n=12-13). Bar graph shows quantification of pair corr. at short distances (5-10 nm). **(c)** Pair corr. analysis of two-color dSTORM imaging labeling GABAAγ2 and GM1 in primary neurons cultured from wildtype and SREBP2 KO animals (n=5-6). Bar graph shows quantification of pair corr. at short distances (0-5 nm). **(d)** Cholesterol levels in ACM compared to 10% FBS DMEM media and neuroblastoma 2a (N2a) cell conditioned media (NCM) (n=3). The neuronal cells produce no cholesterol and serve as a negative control for astrocytes. **(e)** Cholesterol assay on cultures of primary cortical neurons (containing 1% mixed astrocytes) uptake cholesterol from ACM (n=9-15). All data are expressed as mean ± s.e.m., ns, not significant, *p<0.05, **p<0.01, ****p<0.0001, unpaired t-test. B-c each n is a biological replicate. D-e each n is a technical replicate.

Treatment with ACM resulted in a significant increase in the colocalization of GABA_A_R with GM1 clusters by 35% ± 15% (p<0.05) (Fig. 4b), indicating an effect of cholesterol from ACM on receptor clustering. This enhancement was also observed in N2a cells with a 137% ± 50% increase (p<0.05) (Fig. S4a). Given the role of ApoE as the primary lipid transporter in the brain, predominantly for cholesterol^9^, we further explored the effects of lipidated ApoE treatment on GABA_A_R and GM1 colocalization. Both primary neurons and N2a cells showed a similar increase in colocalization upon treatment with lipidated ApoE (Figs. S4b-c).

#### Astrocyte-specific SREBP2 knockout

To contrast the above findings, we examined cultures from astrocyte specific SREBP2 knockout animals, where SREBP2 acts as a key regulator of cholesterol synthesis. The knockout leads to a reduction in cholesterol production, affecting the primary source of cholesterol for neurons^9,58^. Cultures derived from these knockout animals exhibited a significant decrease (75% ± 17%, p<0.01) in the colocalization of GABA_A_R and GM1 clusters (Fig. 4c), underscoring the importance of astrocyte-derived cholesterol in receptor localization.

#### ACM treatment

To validate the uptake of cholesterol into cells post-ACM treatment, we measured the cholesterol levels in ACM and in cells after treatment. The cholesterol concentration in ACM was found to be approximately threefold higher than that in standard 10% FBS DMEM media (Fig. 4d). As a control, N2a conditioned media (NCM) was compared, showing a significant reduction in media cholesterol content after culturing in 10% FBS DMEM media for 3 days (20% ± 5%, p<0.05). ACM treatment significantly increased cholesterol levels in primary neurons (8% ± 1%, p<0.0001), whereas NCM treatment did not alter cholesterol levels (Fig. 4e).

The effect of ACM on cholesterol loading was also assessed in N2a cultures (Fig. S4d), where ACM treatment led to a significant increase in cholesterol levels (15% ± 2%, p<0.0001). Interestingly, NCM treatment resulted in a slight increase in cholesterol levels (7% ± 3%, p<0.05), suggesting that signaling molecules present in NCM contributed to this effect. In contrast, the treatment with endogenous cholesterol transporter protein ApoE increased serum cholesterol levels in N2a cells but not in primary neurons with 1% astrocytes^9^ (Fig. S4e-f).

### Effects of ACM on GABA_A_R currents in WSS-1 cells

Building on the observation that ACM treatment shifts GABA_A_R to cholesterol rich environment, we aimed to determine whether ACM treatment correlate with a functional change in the receptor’s ion conduction using whole-cell patch-clamp recordings in WSS-1 cells, which overexpress GABA_A_R. To correlate the data with the dSTORM imaging, we used the same treatment, 1 mM GABA and applied it continuously at 3 ml/min for ~10 min. As mentioned, GABA_A_R’s rate of desensitization decreases with increasing concentration of GABA^59,60^.

Figure 5a shows representative traces of GABA_A_R currents elicited with 1 mM GABA in cells with and without overnight treatment with ACM. We found both the time to max current (I_max_) and the time to desensitization were increased significantly (from 12s to 24s, n=6, p<0.001; and from 50s to 75s, n=6, p<0.05 respectively, Fig. 5b-c) after treatment with ACM. Despite the pronounced delay in activation, the magnitude of the I_max_ did not decrease; rather, it appears to increase slightly (Fig. 5d). Combined, these data suggest that for a GABA_A_R activation sequence to happen, the receptor may need to leave the lipid rafts.

**Figure 5.**
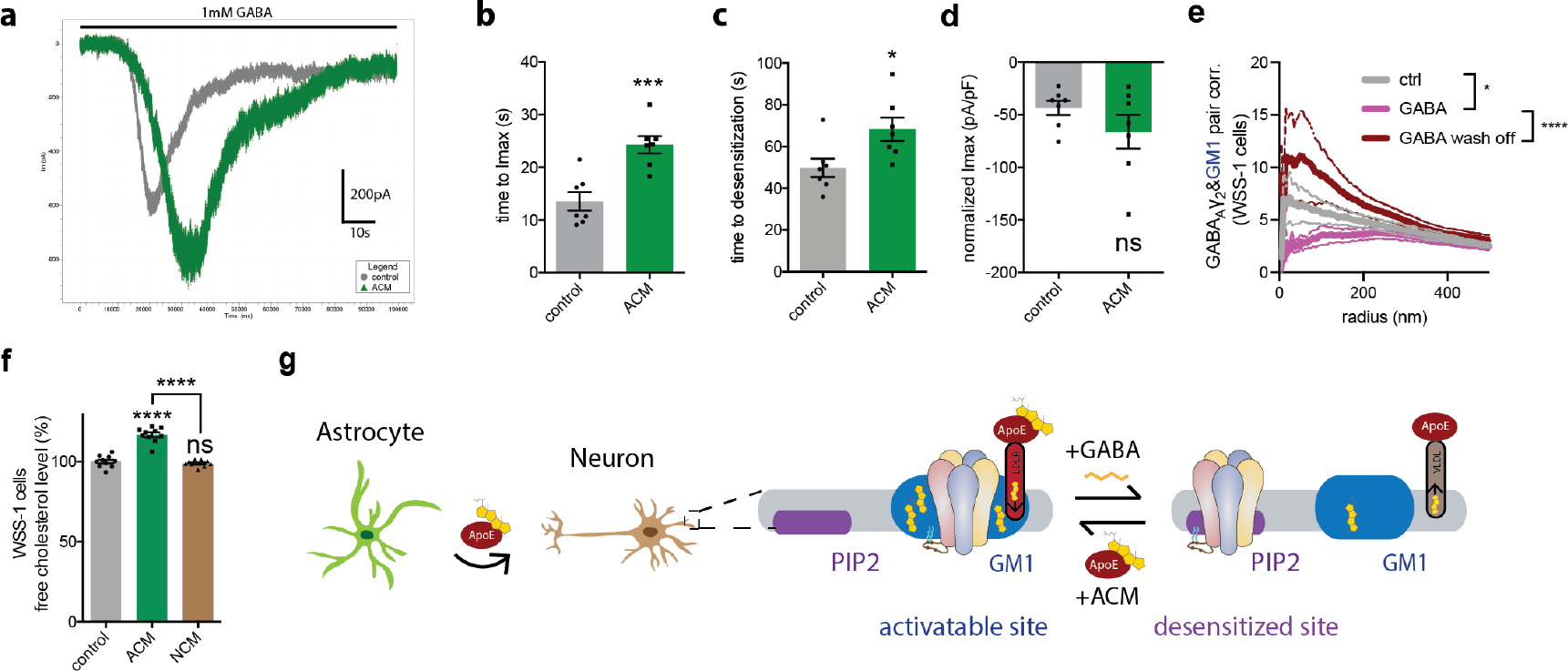
Astrocyte conditioned media delays GABA_A_R membrane desensitization. **(a)** Two examples of raw traces of whole-cell patch recording. The bar indicates continuous application of 1mM GABA. A control cell is shown in grey, and an ACM treated cell is shown in green. **(b-d)** The time to reach (b) maximum current (Imax), (c) desensitization and (d) the normalized maximum current of whole-cell patching on WSS-1 cells incubated with astrocyte conditioned media (ACM) (n=7). **(e)** GABA_A_γ2 /GM1 cross-pair correlation (pair corr.) analysis of WSS-1 cells treated with 1 mM GABA for 10min and after GABA wash off and recovery for 15min (n=6-7). The statistical comparison of the curves is analyzed at 10-30 nm radius with nested t-test. **(f)** Cholesterol assay on WSS-1 cells with and without uptake of astrocyte-derived cholesterol (n=10). **(g)** A cartoon diagram of depicting uptake of cholesterol into neurons from ACM via apolipoprotein E (ApoE) containing lipid particles and its receptor low density lipoprotein receptor (LDLR). The astrocyte cholesterol recruits GABA_A_γ2 to GM1 clusters (blue shading). Prior to GABA treatment, the receptors are bound to GM1 clusters in the activatable state. As part of the activation process, GABA_A_R leaves GM1 lipids and associates with PIP_2_ clusters. All data are expressed as mean ± s.e.m., ns, not significant, *p<0.05, ***p<0.001, ****p<0.0001, unpaired t-test. Each n is a biological replicate, except in (f) which are technical replicates.

To confirm that GABA activation is dependent on GABA_A_R lipid environment the same in WSS-1 cells as they do in primary neurons (Figs. 1-2), we performed dSTORM analysis and cross-pair correlation of GABA_A_R with GM1 in WSS-1 cells. Consistent with primary neurons, GABA decreased the association of GABA_A_R with GM1 lipids in WSS-1 cells and the effect was reversible. And like primary neurons, upon reversal the GABA_A_R/GM1 cross-pair correlation was increased compared to starting conditions. Hence, these data show that the regulation of the GABA_A_R lipid environment extends to diverse cell types.

Additionally, to confirm cholesterol uptake, we tested cholesterol levels in WSS-1 cells after ACM treatment. Like uptake into primary neurons, ACM treatment increased the free cholesterol levels within WSS-1 cells significantly (~20%, Fig. 5f), further suggesting cholesterol is at least one component of ACM regulating the channel through paracrine like signaling.

Lastly, to confirm that GABA_A_Rs were on the surface during our electrophysiology experiments, we measured GABA_A_R protein levels with and without ACM treatment using fluorescent immunostaining (Fig. S5a-b). This is an important experiment since the movement of GABA_A_R to GM1 lipids could affect endocytosis^27^, a second known mechanism of GABA_A_R desensitization^61^, which we do see in primary neurons as discussed below. However, in WSS-1 cells we saw no significant change in GABA_A_R protein levels on the surface after ACM treatment in WSS-1 cells. The reason is unclear, but it confirms that GABA_A_R can desensitize on the surface independent of receptor mediated endocytosis.

### Astrocytic regulation of GABA_A_R endocytosis

As mentioned, GM1 clusters serve as pivotal sites for GABA_A_R internalization, facilitating clathrin-mediated and dynamin-dependent endocytosis^62,63^. Endocytosis is also a well-established mechanism of GABA_A_R desensitization separate from conformation mediated desensitization. It works through removal of receptors from the membrane where the channel functions. Given our findings that astrocyte-derived cholesterol influences GABA_A_R localization to GM1 clusters, we further investigated how ACM treatment affects GABA_A_R endocytosis. In both primary neurons and N2a cells, ACM treatment reduced GABA_A_R’s membrane expression by 12% ± 3% in primary neurons and 8% ± 1% in N2a cells without altering overall cellular expression levels (Fig. S6a-d).

#### Methyl-β-cyclodextrin (MβCD) vs. Avasimibe

To delineate the influence of cholesterol on GABA_A_R regulation, we employed endocytosis inhibitors that alter membrane cholesterol levels in N2a cells. First, we used methyl-β-cyclodextrin (MβCD), an endocytic blocker^64^, to acutely remove membrane cholesterol, disrupting GM1 clusters. MβCD treatment reduced the cross-pair correlation of GABA_A_R with GM1 clusters while increasing its association with PIP_2_ clusters, suggesting that MβCD facilitates receptor release and relocation to PIP_2_ clusters upon cholesterol removal (Fig. S6e-f). Conversely, avasimibe (AVS), an inhibitor of Acyl-CoA: cholesterol acyltransferase (ACAT) that blocks cholesterol internalization, while raising cholesterol levels on the membrane^58^, increased GABA_A_R’s cross-pair correlation with GM1 clusters and decreased its association with PIP_2_ clusters. This result shows AVS promotes GABA_A_R recruitment to GM1 clusters, supporting the finding that AVS reduces endocytosis by trapping receptors in GM1 clusters on the surface. These data show that when endocytosis is blocked, the movement of GABA_A_R between GM1 and PIP_2_ clusters follows cholesterol’s change.

#### Phospholipase D

The role of PLD, a necessary enzyme for endocytosis^65,66^, in GABA_A_R endocytosis was explored by comparing cells transfected with catalytically inactive PLD (xPLD) to those with active mouse PLD (mPLD). xPLD transfection slightly reduced cholesterol levels and significantly increased both the surface and overall expression levels of GABA_A_Rs, suggesting an inhibition of receptor endocytosis (Fig. S5c-e). In contrast, mPLD transfection had no impact on overall expression but decreased membrane expression, aligning with PLD’s role in facilitating endocytosis. Whole-cell patch-clamp recordings in xPLD-transfected WSS-1 cells showed no change in maximal current (Fig. S5f) but a significant extension in desensitization time (65% +/- 14% (p<0.01)) (Fig. S6g). This data further supports that GABA_A_R can desensitize locally on the membrane.

Figure S6h illustrates a cellular model to summarize the two distinct desensitization mechanisms of GABA_A_R as endocytic desensitization (receptor removal) and membrane desensitization (conformational change). In the endocytic desensitization, high cholesterol levels resulting from ACM treatment directs GABA_A_R to GM1 clusters for endocytosis. In the membrane desensitization, GABA_A_R in GM1 clusters are activatable and they traffic to PIP_2_ clusters where they desensitize locally on the membrane.

## Discussion

This study elucidates the critical role of astrocytes and GABA in regulating the lipid environment of GABA_A_R in neurons. GABA initiates a movement of GABA_A_R from cholesterol rich lipids to PIP2 lipids, hence GABA regulates the exposure of GABA_A_R to a lipid environment distinct from the resting/activatable state. Astrocytic secretions have the opposite effect, they move GABA_A_R toward cholesterol rich domains. The lipid environment critically regulates GABA_A_R function^6,67,68^ and this extends to disease^69–72^. Hence understanding the factors that control GABA_A_R’s lipid environment in a cellular membrane is critical for understanding the receptor’s function in the synapse.

Four experiments suggest the movement between lipid environments is physiologically relevant. One, the agonist induces the movement (Fig. 1d). Two, the movement is reversible upon agonist removal (Fig. 2). Three, competitive antagonist blocks the movement (Fig. 1d). And four, the movement correlates with delayed desensitization and increased endocytosis. The association with lipid rafts has also been seen in detergent resistant membranes^73^.

Nonetheless, the mechanisms of GABA and astrocytic regulation remains somewhat vague. Cholesterol is the main component of astrocytic lipid particles^9^. And its uptake could change the concentration sufficient to bind to the receptor. But perhaps more importantly, the receptor moves to a cholesterol rich compartment. This change in cholesterol concentration, based on the location of the receptor could facilitate sufficient cholesterol binding and elicit a conformational change in the protein without dramatically changing the overall concentration of cholesterol in the membrane. Supporting this conclusion, the effects of ACM on GABA_A_R currents occurred with a 10% change in overall cholesterol levels (Fig.4e). This change in concentration alone is unlikely sufficient to saturate previously unoccupied binding sites. However, combined with the movement to a cholesterol rich domain, we expect the concentration of cholesterol the channel experiences locally could increase dramatically and regulate the channel by direct binding^6^.

The movement between GM1 and PIP2 environments also implies a change in hydrophobic thickness and a potential indirect role of lipids in the GABA_A_R’s conformation^74^. When the receptor associates with GM1 clusters, prior to activation, the membrane is thick (~40 Å)^75^. After activation, the receptor associates with PIP_2_ clusters which are reside thin disordered lipids (29-31 Å; see figure S7). These findings suggest the activatable state may have a similar thickness to lipid rafts. Structures solved in nanodisc have an estimated thickness of 31.4 Å. This is likely due to the thickness of the disordered lipids used in the reconstitution. When GABA_A_R associates with GM1 lipids, it would likely need to change conformation to accommodate the increased thickness, for example tilting a helix vertical or removing the bend from a helix. A helix tilting mechanism has been proposed for the nicotinic acetylcholine receptor (nAChR)^70,71^, a cationic homolog of GABA_A_R^72^, and for TREK-1 potassium channels^76,77^. Trapping GABA_A_R in the most accurate activable state may require a hydrophobic thickness closer to 40 Å.

The application of ApoE and ACM to manipulate lipid levels in cell cultures provides key insights into astrocytes regulatory capacity. The increased clustering of GABA_A_R and the diminished clustering in astrocyte specific SREBP2 knockout cultures, suggests a signaling role for cholesterol. We did not knockout apoE here, but previous KO experiments showed apoE is required for the clustering of protein in neurons^9^. The finding is like ACM controlling TREK-1 localization in neurons^76^. However, in those studies, the localization of TREK-1 directly exposed the channel to lipid agonists, whereas with GABA_A_R the nanoscopic change in localization affected desensitization and endocytosis (Fig. 4 and Fig. S6)).

One limitation of the study is the use of neurons without synapses, we relied on gephyrin and GM1 lipids to identify the location most physiologically relevant to synaptic signaling^78^. This strategy helped define the individual roles of the astrocytes relative to the neurons. But similar experiments could be done in the presence of well-defined synapses. A second limitation is that of using multidentate antibodies and toxin during live imaging, this could cause clustering and increase the size of lipid rafts^24,33^. Since we utilized localization, not cluster size, this should not affect our conclusion. Regardless, in our case, the apparent GABA_A_R and GM1 cluster size were smaller live compared to fixed (181.5 nm (Fig.3f) vs 211.7 nm (Fig. 2e) for GABA_A_R, 177.6 nm (Fig.3g) vs 214.5 nm (Fig.2c) for GM1). This suggests the live imaging is not contributing to increased clustering, rather, the fixing processes is likely slower and allows molecules to diffuse and artificially increasing the apparent raft size (~12 nm calculated from the difference in the two radii). For the conclusions of this study, we used the relative location of GABA_A_R in lipid rafts and PIP_2_ domains which are separated by 100-200 nm^27^ (Fig. S1f) and should not be affected by this 12 nm of increased diameter due to fixing. However, it is a consideration, and for systems where the induced clustering is more pronounced or the distance between domains is smaller, these artifacts could be a problem for proper interpretation of the results.

While this study provides foundational insights into cholesterol’s role in GABA_A_R regulation, further research is necessary to fully decipher the molecular details of this regulation and its implications for neuronal physiology and potential therapeutic interventions. Low GABA function has been proposed as a biomarker for mood disorders^79^. Our observation of increased GABA_A_R endocytosis from higher cholesterol levels suggests a potential role of cholesterol and other astrocyte lipids in mood disorders through regulating GABA_A_R function. And Alzheimer’s disease (AD) patients are known to have increased lipids^9^ and this correlates with hyperexcitability of nerves in the AD brain^80^. Additionally, the potential regulatory effect of cholesterol on other ion channels, such as glycine receptors (GlyRs), suggests a broader spectrum of cholesterol-mediated modulation of synaptic signaling.

In conclusion, our study highlights the critical role of astrocyte cholesterol in modulating the lipid environment of GABA_A_R, providing valuable insights into the molecular basis of GABAergic signaling. These findings not only advance our understanding of neuronal signaling mechanisms but also open new avenues for exploring the impact of astrocytic lipids on synaptic function and its potential as a target for therapeutic intervention in neurological disorders.

## Methods

### Cell Culture Procedures

Primary neurons and astrocytes were harvested from embryonic day 17 brain cortices. Primary neurons were cultured in neurobasal medium (gibco, #21103-049) with 2% B27 (gibco, #17504-044) and 1% Glutamax (gibco, #35050-061) at 37 °C with 5% CO_2_. Astrocytes, Neuro-2a (N2a), WSS-1, human embryonic kidney 293T (HEK293T) cells were cultured in Dulbecco’s Modified Eagle Medium (DMEM, Corning™, #10-013-CV) with 10% fetal bovine serum (FBS, Sigma-Aldrich, #F0926) and 1% Penicillin Streptomycin (PS, Corning™, #30-002-Cl) at 37 °C with 5% CO_2_.

Astrocyte conditioned media (ACM) was collected at 3-day intervals from the primary astrocyte culture. All collected ACM was pooled together, divided into aliquots, and then frozen and stored at −20°C for future use.

### dSTORM Imaging Protocol

#### Fixed cells

Cells were initially treated and fixed with 4% paraformaldehyde and 0.1% glutaraldehyde for 15min and reduced with 0.1% sodium borohydride for 7min. For co-labeling the receptors and PIP_2_ lipids, cells were permeabilized with 0.2% Triton X-100 for 15min. Cells intended for co-labeling with GM1 lipids were not permeabilized, and no Triton X-100 was used in the following steps. Cells were blocked with 10% bovine serum albumin (BSA) and 0.05% Triton X-100 in PBS for 90 min. Cells were then labeled with fluorescence-conjugated antibodies for 60min. GABA_A_R was labeled with GABA_A_R γ2 antibody (Novous Biological, NB300-151) conjugated with Atto 647N NHS ester (Sigma-Aldrich, 18373-1MG-F) or Cy3B NHS ester (GE Healthcare, PA63101). GM1 was labeled with cholera toxin subunit B (CTxB) Alexa-647 (Invitrogen, C34778) or CTxB Alexa-555 (Invitrogen, C34776). PIP_2_ was labeled with PIP_2_ antibody (Echelon, Z-P045) conjugated with Cy3B NHS ester or Atto 488 NHS ester (Sigma-Aldrich, 41698-1MG-F). Following the labeling, cells were washed 5 times with 1% BSA and 0.05% Triton X-100 in PBS with 15 min of shaking in-between. The cells were then washed with PBS once for 5 min and post-fixed with the fixative solution for 15 min. Vutara VXL was used for imaging and Vutara SRX software was used for analyzing the imaging data. 3000 frames were collected for each laser channel. Cells were imaged in 10% glucose, 50mM Tris, 10mM NaCl with 1% β-mercaptoethanol, 56 ug/ml glucose oxidase and 34 ug/ml bovine catalase.

*Live cells* were labeled with GABA_A_Rγ2-647 and CtxB-555 for 30min in fresh cultural media then washed with antibody-free fresh cultural media for 3 times, 2min each time. Imaging was performed at 200 fps, repeating 800 frames for GABA_A_Rγ2-647 and 200 frames for CtxB-555. The live cell data were binned every 10s.

### Electrophysiology

Whole-cell patch-clamp recordings of GABA_A_R current were made in WSS-1 cells. Cells were cultured on coverslips. The electrode solution contained 140 mM KCl, 2 mM MgCl_2_, 11 mM EGTA, 0.1 mM ATP, and 10 mM HEPES-KOH (pH 7.4). The extracellular solution contained 140 mM NaCl, 4.7 mM KCl, 1.2 mM MgCl_2_, 2.5 mM CaCl_2_, 11 mM glucose, and 10mM HEPES-NaOH (pH 7.4). The electrode solution had an osmolarity of 300 mOsm and the extracellular solution was at 325 mOsm. Fresh buffer and GABA solution were perfused at 3mL/min. The data was visualized and analyzed by ClampFit.

### In vitro free-cholesterol assay

In vitro free-cholesterol assay was measured in lysed cells and analyzed with an Amplex Red-based assay. Cells were cultured in complete media in 96-well plates until ~90% confluent and lysed with RIPA buffer. The RIPA buffer solubilized the lipids and gives access to cholesterol in all the cellular membranes. The samples were mixed with assay buffer containing 100 μM Amplex red, 2 U/mL horseradish peroxidase (HRP), 4 U/mL cholesterol oxidase, and 4 U/mL cholesteryl esterase in PBS to reach 100 μL of reaction volume. The assay reaction was measured by a fluorescence microplate reader (Tecan Infinite 200 PRO) at an excitation wavelength of 530 nm and an emission wavelength of 585 nm. Relative cholesterol concentration was determined by fluorescence activity and calculated by subtracting the background activity of the reaction buffer without samples.

### In vitro cellular PLD assay

In vitro cellular PLD2 activity was measured in cultured cells by an enzyme-coupled product release assay using amplex red reagent. Cells were cultured in complete media in 96-well plates until ~90% confluent. The PLD reaction was initiated by adding 100 μL of reaction buffer (100 μM amplex red, 2 U/ml HRP, 0.2 U/ml choline oxidase, and 60 μM C8-PC in PBS). The assay reaction was performed for 2h at 37 °C, and the activity was kinetically measured with the fluorescence microplate reader (Tecan Infinite 200 Pro) at an excitation wavelength of 530 nm and an emission wavelength of 585 nm. Relative PLD activity level was determined by fluorescence activity and calculated by subtracting the background activity of the reaction buffer with treatment and without cells.

### Statistical analysis

All statistical analysis were performed in GraphPad Prism 10. For the Student’s *t* test and nested t-test, significance was calculated using a two-tailed, unpaired parametric test with significance defined as \**P* < 0.05, \*\**P* < 0.01, \*\*\**P* < 0.001, and \*\*\*\**P* < 0.0001. All data are expressed as mean ± s.e.m., each n is a biological replicate. Every experiment is repeated at least twice.

## Acknowledgment

We thank Jackson Carter for the help with electrophysiology, Andrew Hansen for the development of Nutmeg software for exporting data and discussion, Julian Bois for the help in maintaining lab cell cultures, Jenna Wingfield and Kaushik Chanda from Sathya Puthanveettil Lab for providing E17 cortical tissues, Michael Froman from Stony Brook for the mouse PLD and mutant PLD cDNA, John Baenziger of University of Ottawa for helpful discussion, and Heather Ferris at the University of Virginia for SREBP2 KO cells. This work was supported by an R01 to S.B.H. (R01NS112534) from the National Institute of Health. We are grateful to the JPB foundation for the purchase of the Vutara VXL microscope. The authors declare no conflict of interest.

---

**Supplemental figure 1.**
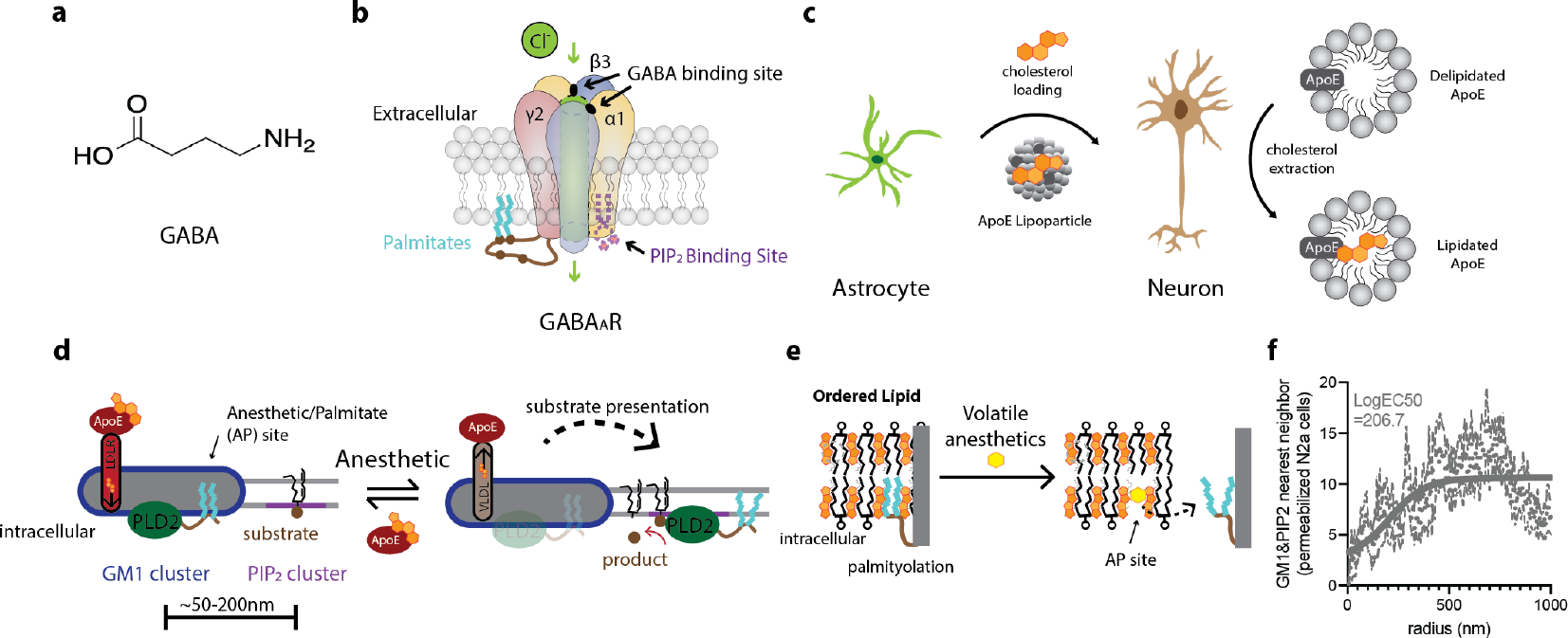
Background for GABA_A_R modulation. **(a)** Molecular structure of GABA. **(b)** A cartoon diagram of *α*1β3γ2 GABA_A_ receptor showing palmitoylation sites at γ2 subunit and PIP_2_ binding sites at *α*1 subunit. **(c)** A cartoon diagram of cholesterol production and trafficking in the brain. Astrocytes are the primary cholesterol suppliers in the brain and astrocyte-derived cholesterol is packed with apolipoprotein E (ApoE) to deliver astrocytic cholesterol to the neuronal membrane. In the absence of astrocytic cholesterol, delipidated apoE extracts cholesterol from the neuronal membrane. **(d)** A cartoon diagram explaining the mechanism of substrate presentation using PLD2 as an example. Palmitate lipids, covalently attached to the PLD2 cause the enzyme to associate with GM1 clusters. Uptake of cholesterol increase the affinity of the palmitates for the GM1 clusters. Anesthetics compete with palmitate within GM1 lipids (the anesthetic/palmitate (AP) site) and this releases the enzyme from GM1 clusters allowing it to diffuse to PIP_2_ clusters where it has access to its substrate. GM1 and PIP_2_ are shown forming distinct clusters on the membrane (~50-200 nm apart). **(e)** A detailed view of the AP site. The palmitates of a protein are shown binding to ordered lipids through favorable lipid-lipid packing. Anesthetics are shown competing for the palmitate site within ordered lipids. **(f)** Average distance (nm) of GM1 clusters and PIP_2_ clusters in N2a cells. The distance is calculated through the half maximal of the Nearest Neighbor analysis in the Vutara software.

**Supplemental figure 2.**
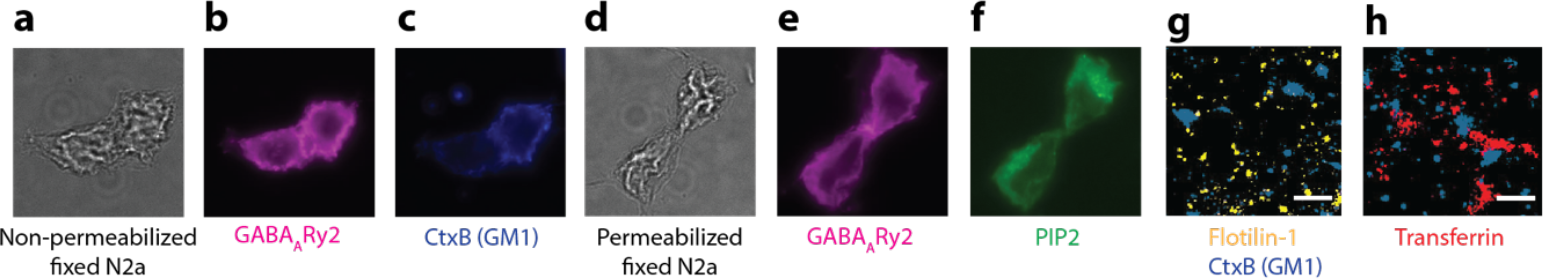
Labeling of fixed N2a cells. **(a)** A trans-luminescent widefield image of non-permeabilized fixed N2a cells. **(b)** A widefield image of fluorescently labeled GABA_A_R in non-permeabilized N2a cells. **(c)** Widefield images of labeled GM1 lipids in non-permeabilized fixed N2a cells. **(d)** A trans luminescent widefield image of permeabilized N2a cells. **(e)** A widefield image of GABA_A_R labeling in permeabilized N2a cells. **(f)** A widefield image of GM1 labeling in permeabilized N2a cells. **(g)** A representative 2-color dSTORM image of Flotilin-1 and GM1 labeling in mouse primary cortical neurons. **(h)** A representative 2-color dSTORM image of Transferrin and GM1 labeling in mouse primary cortical neurons.

**Supplemental figure 3.**
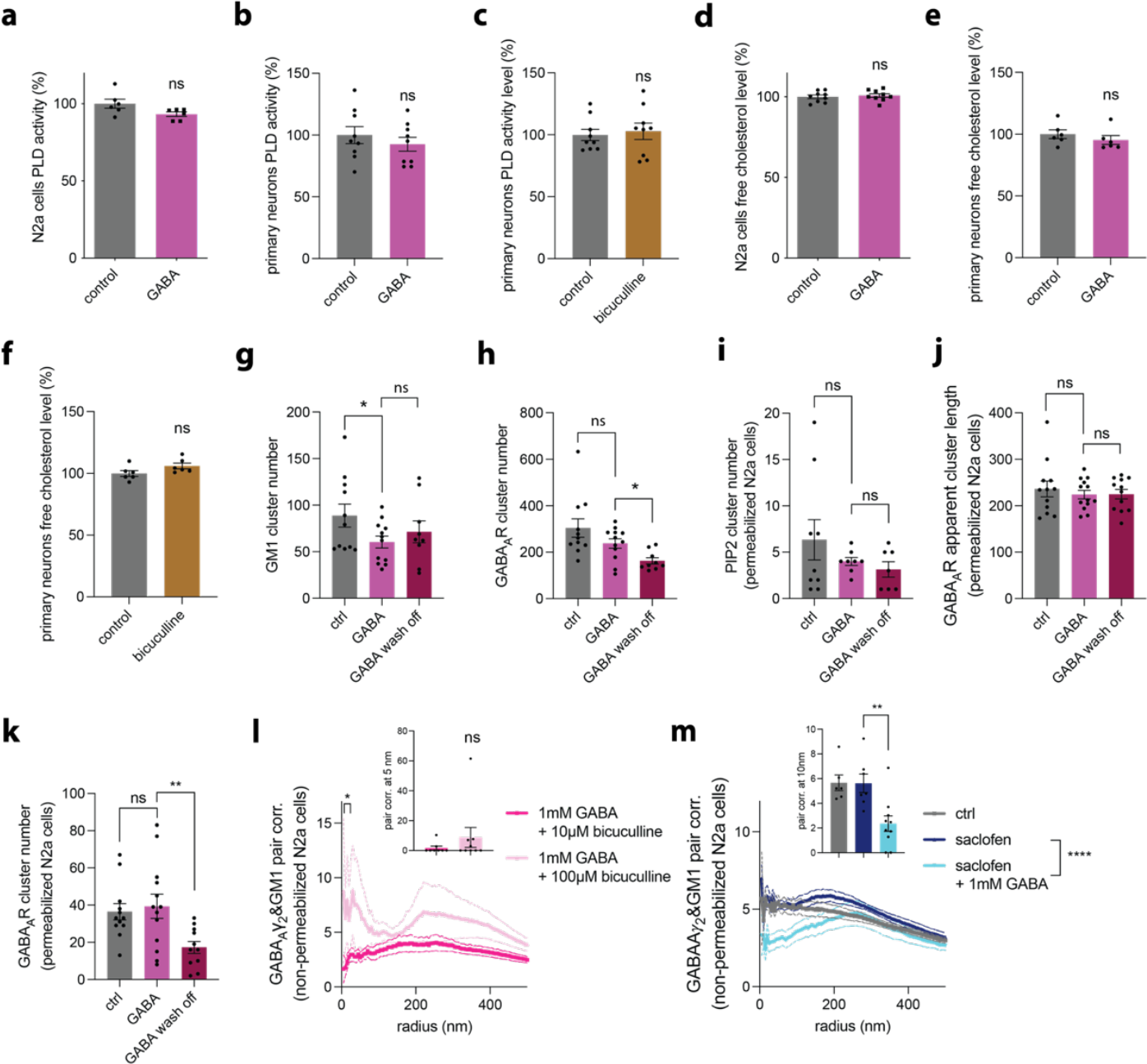
GABA_A_R modulation by agonist and antagonists. **(a)** Quantification of PLD activity in N2a cells treated with 1 mM GABA for 2 h (n=6). **(b)** Quantification of PLD activity in primary neurons treated with 1mM GABA for 2 h (n=9). **(c)** Quantification of PLD activity in primary neurons treated with 10 μM bicuculline for 2 h (n=9). **(d)** Quantification of cholesterol levels in N2a cells treated with 1 mM GABA for 10min. Data are expressed as mean ± s.e.m., ns, not significant, unpaired t-test, n=9. **(e)** Quantification of cholesterol levels in primary neurons treated with 1 mM GABA for 10 mins (n=5). **(f)** Quantification of cholesterol levels in primary neurons treated with 10 μM bicuculline for 10 mins (n=5). **(g)** Number of GM1 clusters in non-permeabilized N2a cells (n=9-12). **(h)** Number of GABA_A_R clusters in non-permeabilized N2a cells (n=9-12). **(i)** Number of PIP_2_ clusters in permeabilized N2a cells (n=7-9). **(j)** Apparent length of GABA_A_R clusters in permeabilized N2a cells (n=12-13). **(k)** Number of GABA_A_R clusters in permeabilized N2a cells (n=12-13). **(l)** Analysis of the cross-pair correlation (pair corr.) of GABA_A_γ2 and GM1 in N2a cells treated with 1 mM GABA together with 10μM bicuculline or 100μM bicuculline for 10min (n=8-9). The statistical comparison of the pair corr. curves is analyzed at radius 5-25 nm with nested t-test, *p<0.05. The bar graph shows quantification at 5 nm radius. **(m)** Analysis of the pair corr. of GABA_A_γ2 and GM1 in N2a cells treated with 100μM saclofen (GABA-derived GABA_B_ antagonist) or 100μM saclofen together with 1mM GABA (n=7-10). The nested t-test is performed at 5-25 nm radius, ****p<0.0001. The bar graph shows quantification at 10 nm radius. All data are expressed as mean ± s.e.m., ns, not significant, *p<0.05, **p<0.01, unpaired t-test. Each n is a technical replicate in (a-f) and a biological replicate in (g-m).

**Supplemental figure 4.**
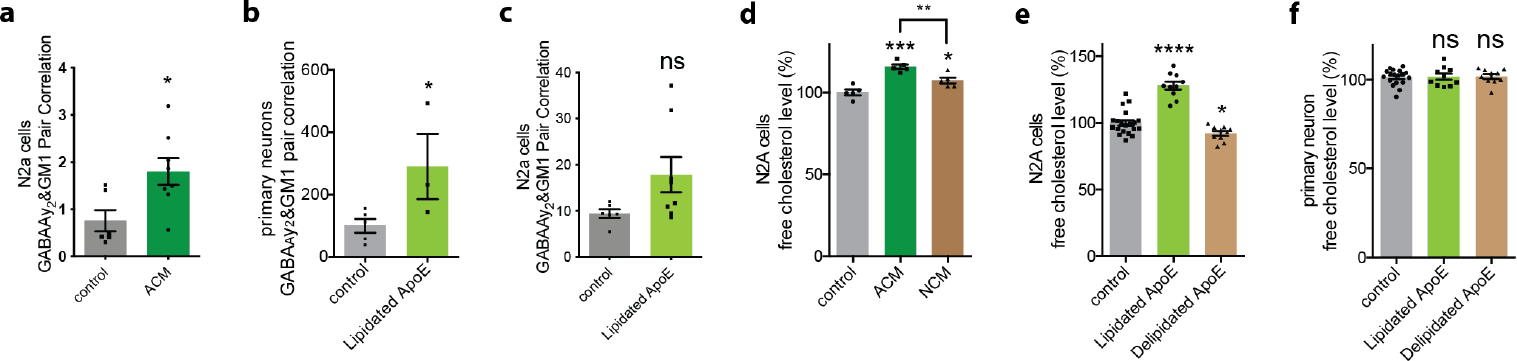
The effects of cholesterol loading on GABA_A_R and the lipid environment. **(a)** Quantification of the pair corr. of GABA_A_γ2 and GM1 at the closest radius in N2a cells with cholesterol loading after treatment with ACM (n=6). **(b)** Quantification of the pair corr. of GABA_A_γ2 and GM1 at the closest radius in primary neurons with cholesterol loading via lipidated ApoE (n=3-5). **(c)** Quantification of the pair corr. of GABA_A_γ2 and GM1 at the closest radius in N2a cells with cholesterol loading via lipidated ApoE (n=6-8). **(d)** Cholesterol assay on overnight cholesterol uptake in N2a cells with ACM and NCM (n=5). **(e)** Cholesterol assay on cholesterol uptake in N2a cells incubated with lipidated (ApoE loaded with 10%FBS DMEM) and delipidated ApoE (ApoE in no FBS DMEM) for 1h (n=10). **(f)** Cholesterol assay on cholesterol uptake in primary neurons (containing 1% astrocytes) with lipidated and delipidated ApoE for 1h. ns, not significant, unpaired t-test, n=10. The percent of astrocytes in our primary neuronal cultures was determined in a previous study^9^. All data are expressed as mean ± s.e.m., ns, not significant, *p<0.05, **p<0.01, ***p<0.001, ****p<0.0001, unpaired t-test. Each n is a biological replicate in (a-c) and a technical replicate in d-f).

**Supplemental figure 5.**
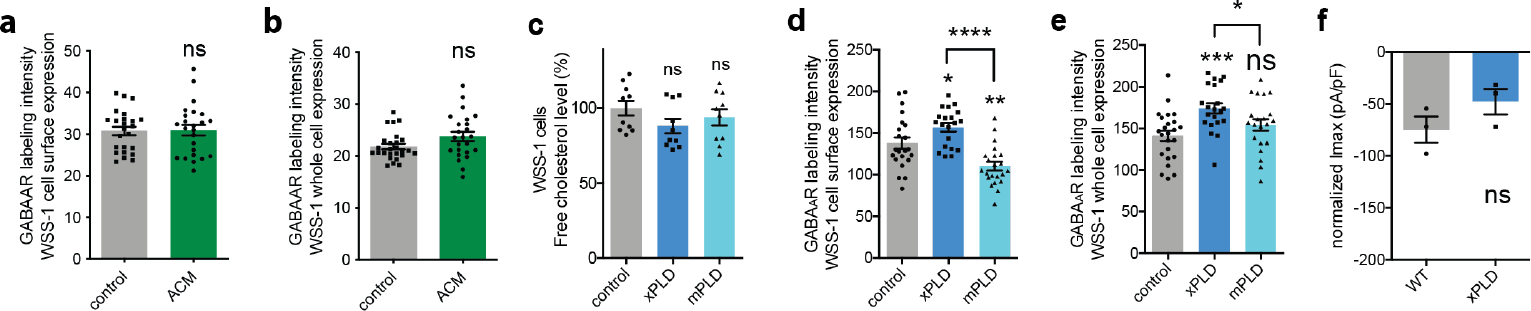
Quantification of GABA_A_R fluorescent labeling in WSS-1 cells. **(a-b)** Quantification of GABA_A_R fluorescent labeling in non-permeabilized (a) and permeabilized (b) in WSS-1 cells indicative of cell surface and whole cell expression respectively (n=24-26). **(c)** Enzymatic assay showing the change in cholesterol levels in WSS-1 cells in the presence of transfected mouse PLD2 (mPLD) and a catalytically inactive variant (K758R, xPLD) (n=10). **(d-e)** Quantification of GABA_A_R labeling intensity in non-permeabilized (a) and permeabilized (b) WSS-1 cells n=20-24 in the presence of over expressed PLDs. **(f)** The normalized maximum whole cell current with and without transfection of xPLD (n=3). All data are expressed as mean ± s.e.m., ns, not significant, *p<0.05, **p<0.01, ***p<0.001, ****p<0.0001.

**Supplemental figure 6.**
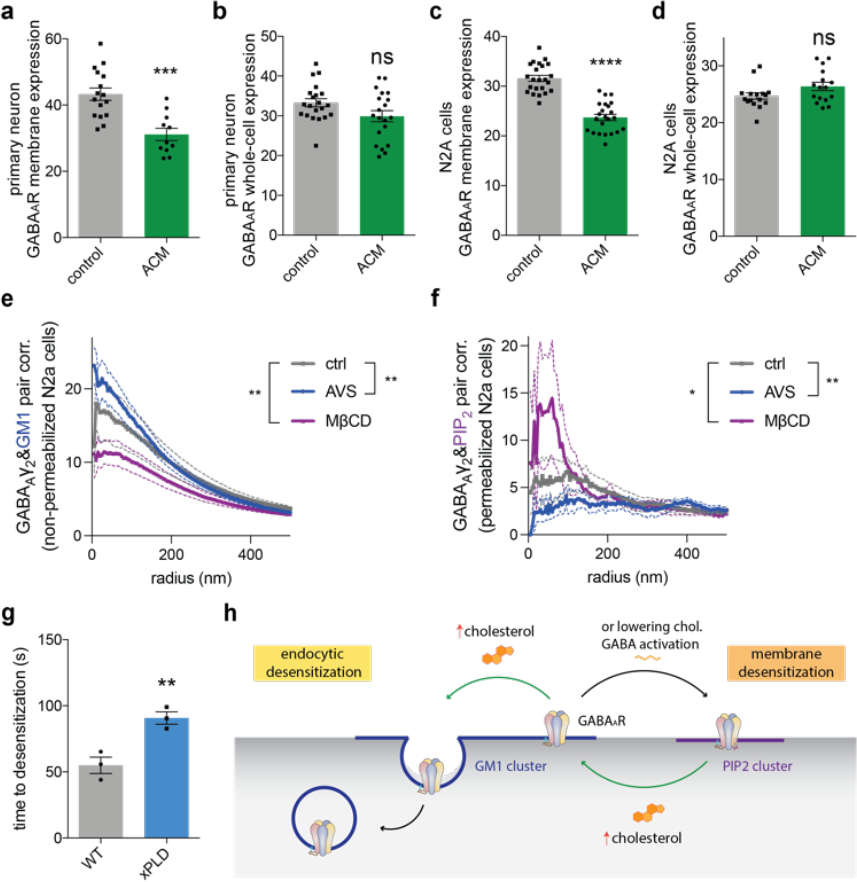
Astrocyte conditioned media induces endocytic desensitization of GABA_A_R in neurons. **(a-b)** Quantification of GABA_A_R surface expression (non-permeabilized, n=11-16) (a) and whole cell (permeabilized, n=20-21) (b) in primary neurons using fluorescent labeling of the receptor. **(c-d)** Quantification of GABA_A_R labeling intensity in N2a cells with (n=23) (c) and without permeabilization (n=16) (d). **(e)** GABA_A_γ2/GM1 cross-pair correlation (pair corr.) analysis with 5 μM avasimide (AVS, an endocytic blocker that works by reducing uptake of cholesterol) or 100 μM MβCD (endocytic blocker that works by disrupting lipid rafts) for 30min in N2a cells. Statistical significance was calculated along the curve (5-30 nm radii) using a nested t-test (n= 6-10). **(f)** The same experiment as in (e) except PIP_2_ was labeled instead of GM1 and the correlation was calculated between GABA_A_γ2 and PIP_2_ (n=4-6) using a nested t-test with radii of 5-35 nm along the curve. **(g)** Shows the effect of transient xPLD expression on GABA_A_R desensitization in WSS-1 cells. The delay in desensitization suggests, in an over expressed system, surface mediated desensitization is slower than desensitization due to endocytosis. **(h)** A cartoon diagram of GABA_A_Rs movement on the membrane. When cells are loaded with cholesterol via ACM treatment (green arrow), GABA_A_Rs move into GM1 clusters where they are internalized through endocytosis. When endocytosis is blocked with AVS or MβCD, the membrane cholesterol regulation effects of these drug traffic GABA_A_Rs into different clusters. All data are expressed as mean ± s.e.m., unpaired t-test, ns, not significant, ***p<0.001, ****p<0.0001, unless otherwise noted.

**Supplemental figure 7.**
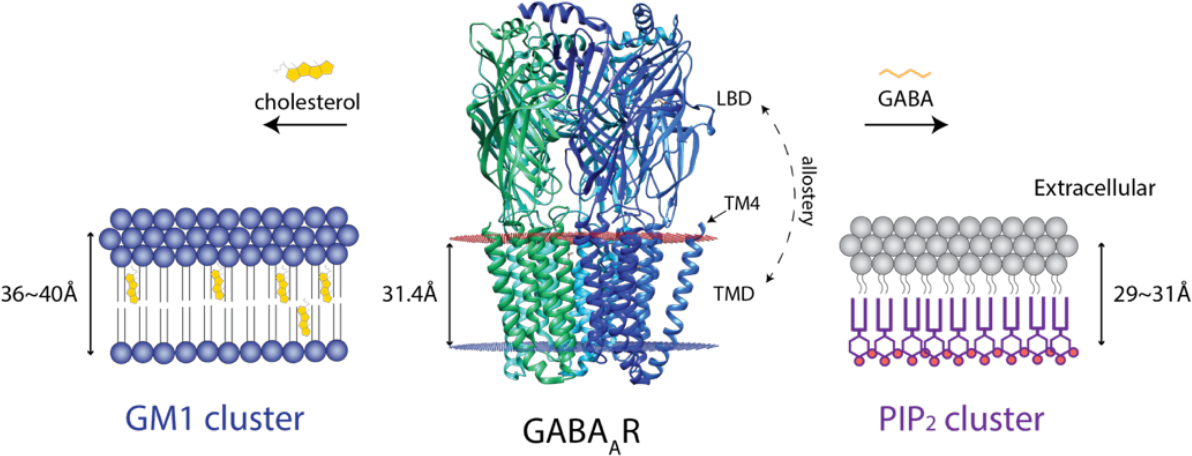
GABA and Astrocytic induced changes in the hydrophobic thickness experienced by GABA_A_R. GABA induces a change in lipid environment from lipid rafts (GM1 clusters, blue shading left) which are ordered and thick (36~40 Å^75^) to PIP_2_ clusters on the inner leaflet (purple shading right) which are thin (29~31 Å). The estimated thicknesses of the lipids are compared to the hydrophobic thickness of the GABAAR structure (31.4 Å, PDB: 6X3T) based on orientations of proteins in membranes (OPM) database (the red shaded surface indicates the extracellular hydrophobic boarder of the membrane and the blue shading, the intracellular boarder calculated from the proteins transmembrane domain (TMD), center panel). The dashed line indicates the allosteric coupling between ligand binding domain (LBD), the site of agonist binding, and the TMD, where the channel gate is located within the membrane. The 31.4 Å hydrophobic thickness is most like PIP_2_ clusters and likely reflects the hydrophobic thickness of the unsaturated lipids used to determine the structure in nanodisc absent cholesterol. When the channel moves to lipid clusters, the transmembrane 4 (TM4) helix in GABA_A_R is likely exposed to a change in hydrophobic thickness. If the TMD were to experience no conformational change in hydrophobic thickness, we anticipate a hydrophobic mismatch of 6-8 Å, upon movement into lipid rafts. TM4 in the homologous nicotinic acetylcholine receptor is thought to tilt with thickness and allosterically couple this tilt through the conserved cys-loop (not shown) ^81^.

